# Changing Times: Fluorescence-lifetime Analysis of Amyloidogenic SF-IAPP Fusion Protein

**DOI:** 10.1101/317917

**Authors:** Olga I Antimonova, Dmitry V Lebedev, Yana A Zabrodskaya, Natalia A Grudinina, Michael M Shavlovsky, Vladimir V Egorov

**Affiliations:** Department of Molecular Genetics, Federal State Budgetary Scientific Institution “Institute of Experimental Medicine”, 197376, Akademika Pavlova St. 12, St. Petersburg, Russia; Department of Molecular and Radiation Biophysics, Petersburg Nuclear Physics Institute named by B. P. Konstantinov of National Research Center “Kurchatov Institute”, 188300, mkr. Orlova roshcha 1, Gatchina, Russia; Department of Molecular biology of Viruses, Research Institute of Influenza, Ministry of Healthcare of the Russian Federation, 197376, Prof. Popov St. 15/17, St. Petersburg, Russia

**Keywords:** FLIM, green fluorescent protein, IAPP, amyloid-like fibrils, atomic force microscopy

## Abstract

The fluorescence lifetime of the superfolder green fluorescent protein (SF) and the SF protein fused with islet amyloid polypeptide (SF-IAPP) were studied in polyacrylamide gel. It was shown that the SF average fluorescence lifetime under these conditions slightly differs from that of the SF-IAPP monomer. SF-IAPP does not lose the ability to form amyloid-like fibrils; meanwhile, the average fluorescence lifetime of the fusion protein in fibrils is reduced. We propose the application of Fluorescent-lifetime Imaging Microscopy (FLIM) to the measurement of average fluorescence lifetimes of fusion proteins (amyloidogenic protein–SF) in the context of studies using cellular models of conformational diseases.

## Introduction

In a number of conformational diseases, for example Parkinson’s disease [1] and type 2 diabetes [2], intracellular accumulation of proteins bearing non-native conformations occurs. The search for compounds that are capable of penetration into the cell, yet also capable of hindering the formation and accumulation of toxic protein aggregates, is an urgent task. It is possible to detect a therapeutic agent’s effect on the intracellular accumulation of amyloid-like aggregates using fluorescent dyes. However, those methods are used after cell fixation, thereby preventing real-time observations. Following the formation of fibrils, a change in the fluorescence lifetime of the dyes covalently bound to the protein monomers is observed [3].

Moreover, it is known that when fibrils are formed, the intrinsic fluorescence of amyloid-like aggregates appears. This is characterized by a specific lifetime [4], and the fluorescence lifetime of aromatic residues changes [5]. Also, it has been shown [6] that in amyloid-like fibrils, the fluorescence lifetime of yellow fluorescent protein fused with alpha-sinuclein decreases. In this study, we analyzed the fluorescence lifetime of superfolder green fluorescent protein (SF) and SF protein fused with islet amyloid polypeptide (SF-IAPP), in both the monomeric and amyloid-like states (when formed).

## Methods

### Recombinant protein

A protein expression construct encoding the SF-IAPP protein was designed using a previously constructed plasmid (based on the commercial pTrc99A_P7 vector) into which the SF sequence was introduced. DNA of the mature IAPP gene, necessary for incorporation into the plasmid, was obtained by PCR using a cDNA library containing the mature IAPP sequence as a template. A forward primer corresponding to the 5’-end of the human mature IAPP coding sequence (5’- cag**gtcgac**<u>cctaggggc</u>aaatgcaacact-3’), containing a Sal I restriction site (**bold**) and a thrombin cleavage site coding sequence (<u>underlined</u>) immediately before the first amino acid residue of IAPP, was used. A reverse primer, corresponding to the 3’-end of IAPP coding sequence (5’- gtc**aagctt***tca*<u>gtggtggtggtggtg</u>atatgtattggatcccac-3’), was designed to contain a Hind III restriction site (**bold**), a stop codon inserted in-frame after the last residue (*italic*), and a sequence which encodes five histidines (<u>underlined</u>) immediately after the last amino acid residue of IAPP.

PCR was performed on a TerCyc amplifier using AmpliSens PCR reagents. Following an initial extended denaturation (of 94°C for 5 min.), PCR consisted of 30 cycles. Each cycle was as follows: at 94°C for 1 min.; 55°C for 1 min.; 72°C for 1 min. An additional final extension was performed at 72°C for 9 min., and ligated into the vector. In order to confirm the presence of the target nucleotide sequence following cloning, the plasmid was sequenced using an M13R standard primer (−48) (AGCGGATAACAATTTCACAC). The sequencing procedure was performed at the Beagle LLC (St. Petersburg, http://www.biobeagle.com/). Table 1 and 2 show the SF-IAPP and SF recombinant protein sequences, as obtained by *in silico* translation, respectively. *Italic* indicates the sequence corresponding to the SF protein; IAPP is <u>underlined</u>.

**Table 1.**
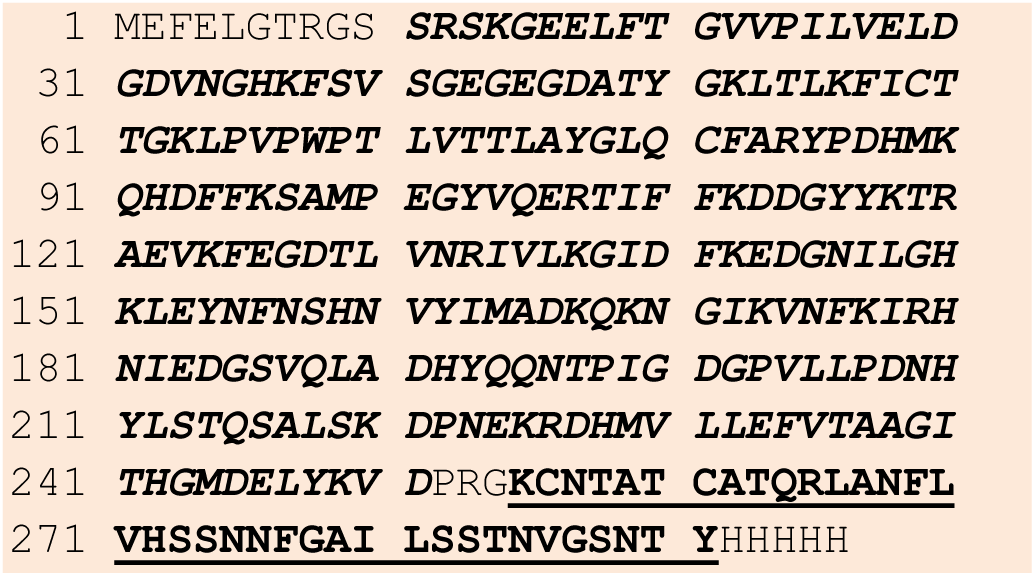
Amino acid sequence of SF-IAPP recombinant protein.

**Table 2.**
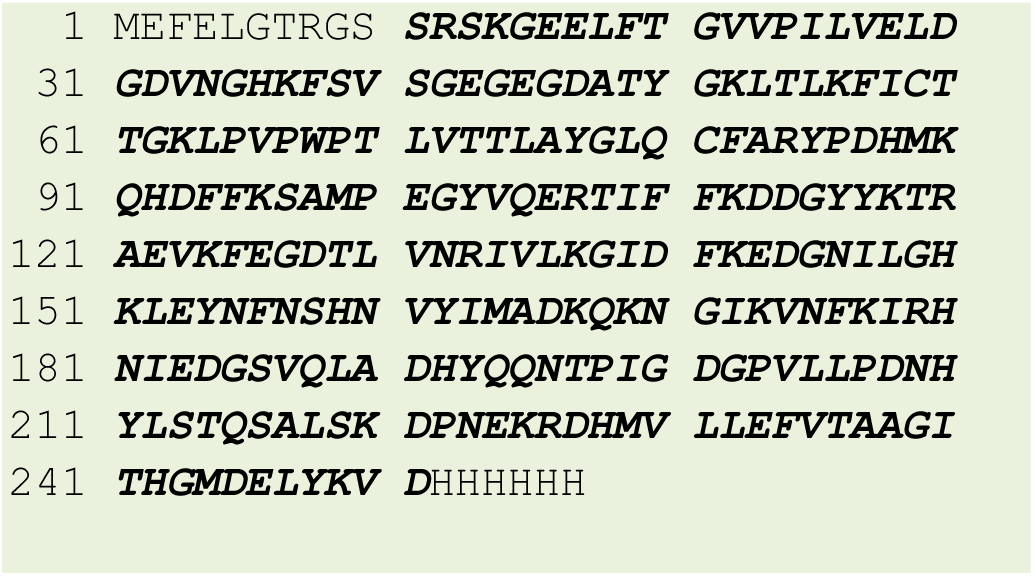
Amino acid sequence of SF recombinant protein.

### Synthesis and isolation of SF-IAPP protein

The SF-IAPP protein was produced in the *E. coli* BL21DE3 expression strain by transformation with the aforementioned plasmid using previously described standard procedures [7]. The concentration of the purified protein was measured spectrophotometrically at the absorption wavelength of the green fluorescent protein (490 nm) using a coefficient E_0.1%_=1.4 for SF и E_0.1%_=1.6 for SF-IAPP.

### Electrophoretic separation of proteins under native conditions

Electrophoretic analysis, under non-denaturing conditions, was carried out according to a standard polyacrylamide gel procedure using a BioRad device (USA). The gel utilized a 5% acrylamide (0.083 M Tris-HCl pH 6.8) stacking portion and a 10% acrylamide (0.75 M Tris-HCl pH 9.2) separating portion. Samples were applied in a buffer containing 0.1 M Tris-HCl pH 6.8, 50% glycerol, and 0.05% bromophenol blue. Tris-glycine buffer pH 8.3 (16.5 mM Tris, 128 mM glycine) was used as the running buffer.

### Fibril production

The SF-IAPP fusion protein was dissolved in a 20mM sodium phosphate buffer (pH 8.0) containing 300mM NaCl and 200 mM imidazole at a concentration of 0.5 mg/ml. 2,2,2-TFE was added to the dissolved protein solution until the concentration of 2,2,2-TFE was 10%.

The resulting solution was incubated overnight in an Eppendorf orbital shaker at room temperature using a 500 rpm rotation speed. The resulting precipitate, obtained by incubation, was examined by atomic force microscopy and by electrophoretic separation under native conditions. A second type of fibril solution was prepared; the manner of preparation was identical to the first, but with the use of mercaptoethanol (100 mM) instead of 2,2,2-TFE.

### Atomic force Microscopy

A suspension of SF-IAPP fibrils (0.13 mg/ml) was placed onto a freshly cleaved mica surface. After 1 min. of incubation at room temperature, the mica surface was washed three times with water to remove buffer constituents. The sample topography measurements were performed in semi-contact mode on an atomic force microscope (NT-MDT – Smena B) using an NSG03 probe (NT-MDT, Russia). The images were analyzed with the assistance of Gwyddion software [8].

### Mass spectrometry

After the electrophoretic separation of proteins under native conditions, the polyacrylamide gel was Coomassie stained [9]. Images were obtained using a digital image station (ChemiDoc XRS+, BioRad). Prior to mass spectrometry, the protein bands of interest were excised and prepared for enzymatic hydrolysis by porcine trypsin (Promega), as described [10]. Next, trypsin was added to the dried gel fragments (20 μg/ml in 50 mM bicarbonate buffer). Enzymatic hydrolysis was performed at 60°C for 2 hours. The reaction was stopped by the addition of an aqueous solution containing 1% trifluoroacetic acid and 10% acetonitrile. The mass spectra of the tryptic peptides were obtained via an UltrafleXtreme MALDI-TOF/TOF mass spectrometer (Bruker, Germany) in positive ion detection mode using the HCCA matrix (alpha-cyano-4-hydroxycinnamic acid, Bruker). For each spectrum, 3000 laser pulses were summed.

Recombinant protein sequences were added to the local database. The identification of proteins was carried out using the Mascot search algorithm (matrixscience.com) with simultaneous access to the SwissProt database and the local database. As possible modifications of the peptides, oxidation (M), deamidation (N), and carbamidomethylation (C) were indicated. The maximum mass error was limited to 20 ppm tolerance.

### FLIM

FLIM measurements were performed with a LEICA fluorescent laser scanning microscope in two-photon excitation mode with a multi-photon laser at 800 nm. The signal was registered using ultrafast detectors. The initial data processing was performed with the SymPhoTime program. The images were taken at a resolution of 256 × 256 pixels, and photon registration was completed when 1,000 photon counts were accumulated. The data obtained was subjected to a fluorescence time analysis in the range of 1.0-12.0 ns. Measurements were carried out on at least three different sections of the band (excised from the native, unstained gel). The numerical values given correspond to the averaging of all measurements from two independent protein preparations per sample.

The spectra were analyzed using Origin2015 software; baseline subtractions were performed. Fluorescence average lifetimes *t (ns)* were calculated for *t*_0_ ≤ *t_i_* ≤ *t_n_* as follow:

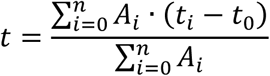

*A_i_* – number of photon counts at *t_i_*,
*t_0_* – time (ns), corresponding to *A_i_* maximum

The resulting spectra were also curved fitted using a timeframe of 1.2-6.2 ns. The lifetimes were calculated by single-exponential (*t1,* ns) or two-exponential (*t1* and *t2, ns*) analysis. The differences between experimental spectra and curve fits were presented in the form of residuals.

## Results and discussion

After the electrophoretic separation of SF-IAPP and SF under native conditions, several bands were observed by means of green fluorescence (Figure 1. lanes 1 and 2, respectively). In lane 1, two bands were observed. It has been hypothesized that the presence of two bands is associated with a change in the electrophoretic mobility of a portion of the SF-IAPP monomers due to deamidation. A polyacrylamide gel, identical to the one shown in Figure 1, was Coomassie stained and the bands analyzed by mass spectrometry in order to confirm the amino acid sequences of the recombinant proteins and in order to test the deamidation hypothesis.

**Figure 1.**
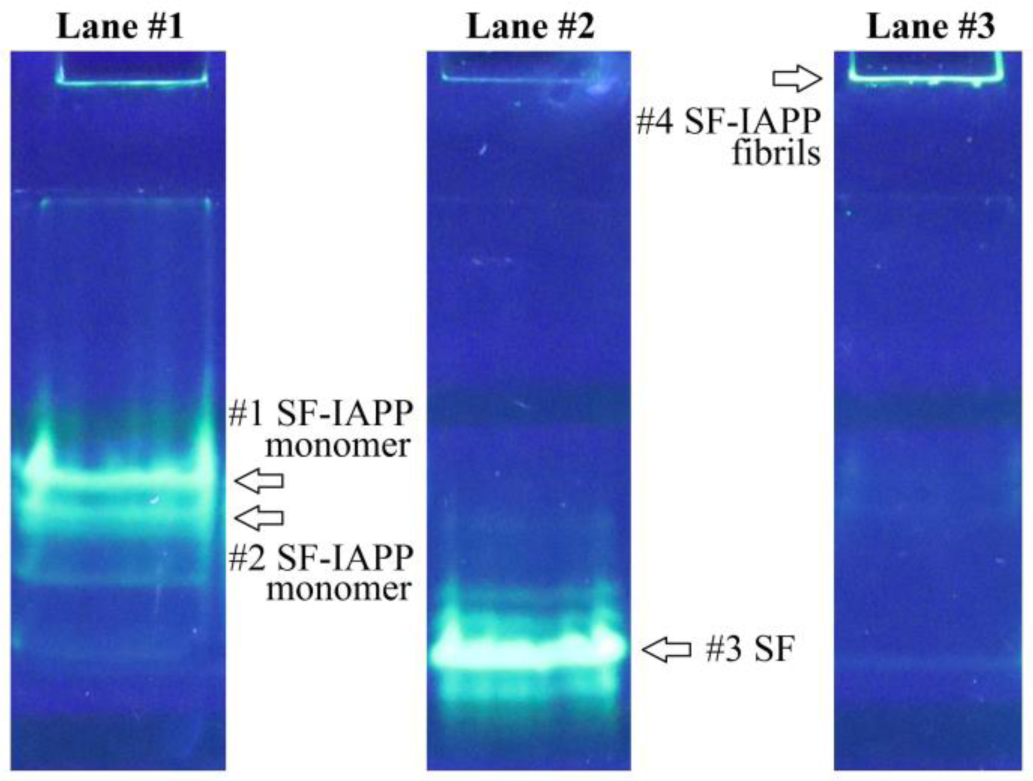
Results of gel electrophoresis under native conditions. The arrows indicate the areas cut out from the sister gel for mass-spectrometric identification. Lanes: 1 SF-IAPP; 2 SF; 3 SF-IAPP fibrils. Locations: #1 and #2 (SF-IAPP recombinant protein); #3 (SF), #4 (SF-IAPP fibrils).

In the band cut from lane 3, the recombinant SF protein (Score 183 at a threshold value of 70) was reliably identified. In bands #1 and #2, the recombinant SF-IAPP protein was reliably identified (Score values were 235 and 231, respectively, at a threshold value of 70). The spectra revealed ions corresponding to the characteristic peptide LANFLVHSSNNFGAILSSTNVGSNTYHHHHH (266-296 a.a.r. fusion protein, Table 1, corresponding to 12-37 a.a.r. IAPP with the His-tag sequence at the C-terminus) with one deamidated residue in SF-IAPP band #1 and two deamidated residues in SF-IAPP band #2. The deamidation of IAPP residues 14 and 21 in the protein and its role in fibrillogenesis is shown [11]. Changing the mass and charge of the protein during deamidation lead to a difference in electrophoretic mobility under native conditions.

Thus, in lane 1, the recombinant SF-IAPP protein was reliably identified. The presence of two bands with different electrophoretic mobilities is associated with a different number of deamidated residues in the C-terminal region of the protein (one in the upper band and two in the lower band).

Measurement of SF and SF-IAPP fluorescence lifetime was carried out after electrophoretic separation of proteins under native conditions (Figure 1). The fluorescence lifetime of the green fluorescent protein is very sensitive to external conditions, such as the ionic strength or the viscosity of the solvent [12]. The use of electrophoretic separation in an *in vitro* system allows one to study the proteins’ fluorescence lifetimes in the same environment, namely when they are incorporated into a polyacrylamide gel. In that immobile state, SF and SF-IAPP samples are in monomeric form, unlike when in solution (in which they can form oligomers differing in fluorescent properties). Fibrils formed from SF-IAPP protein and characterized earlier by atomic force microscopy (Figure 2), were also applied to the gel on lane #3 for comparison purposes (Figure 1).

**Figure 2.**
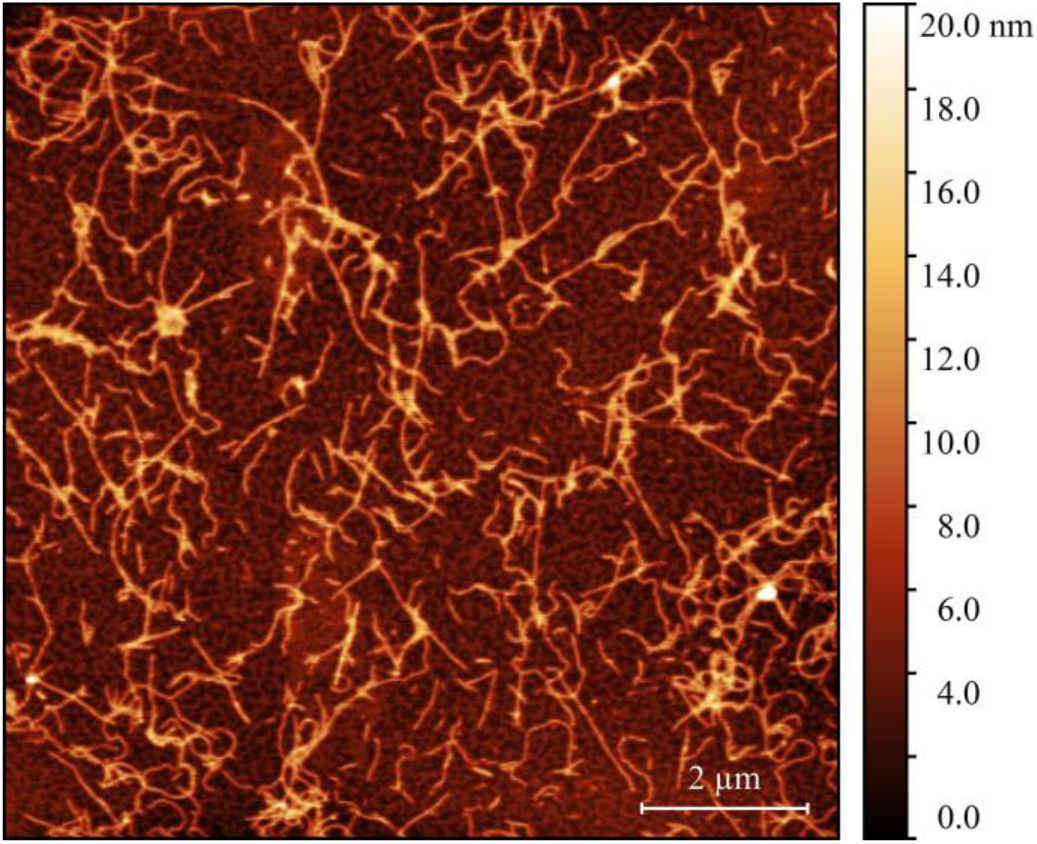
Surface topography of a sample containing SF-IAPP protein fibrils, obtained by atomic force microscopy. The scale bar is 2 μm. On the right is the pseudocolor ruler indicating the particles’ height (nm).

In preparation for FLIM analysis, the excised band fragments indicated in Figure 1 were cut from the gel, placed in buffer, and mounted between microscope slides and their cover slips. The average fluorescence lifetimes were calculated on the basis of four independent measurements and are presented in Table 3. The average fluorescence lifetimes of monomers with one or two deamidations (Figure 1, band #1 and #2) did not differ. Thus, these two forms are indicated with only one term, “SF-IAPP monomer”. Fibrils prepared with or without TFE are presented separately.

**Table 3.** Average fluorescence lifetimes, ns.

|  |  |
| --- | --- |
| SF-IAPP monomer | $2.28 \pm 0.14$ ns |
| SF | $2.51 \pm 0.14$ ns |
| SF-IAPP fibrils | $1.47 \pm 0.21$ ns |
| SF-IAPP fibrils / TFE | $1.89 \pm 0.02$ ns |

Slight differences in the fluorescence lifetime of SF and SF-IAPP protein monomers were observed, while the fluorescence lifetime of SF-IAPP when incorporated into fibrils significantly decreased. Since all of the proteins studied were observed under the same conditions, it can be assumed that the change in the temporal parameters of fluorescence is due to the inclusion of the proteins into amyloid-like aggregates. The almost identical fluorescence lifetimes of the SF and SF-IAPP proteins support this idea. Figure 3a,b represents the single exponential best curve fits (dotted dashed line) and shows residuals below as small circles. It should be noted that all experimental data for SF-IAPP monomers and SF could be approximated by a single-exponential distribution (Figure 3a,b); furthermore, the constant *t1* is in agreement with the fluorescence lifetimes (Table 3). However, for SF-IAPP fibrils, there was a greater discrepancy between the experimental and theoretical curves (Figure 3c, residuals, shown as a line) than for protein monomers in this approximation. Due to this fact, the experimental curve corresponding to SF-IAPP fibrils’ fluorescence decay was approximated by a two-exponential distribution (Figure 3c, best double exponential fit shown as dotted dashed line; residuals - circles).

**Figure 3.**
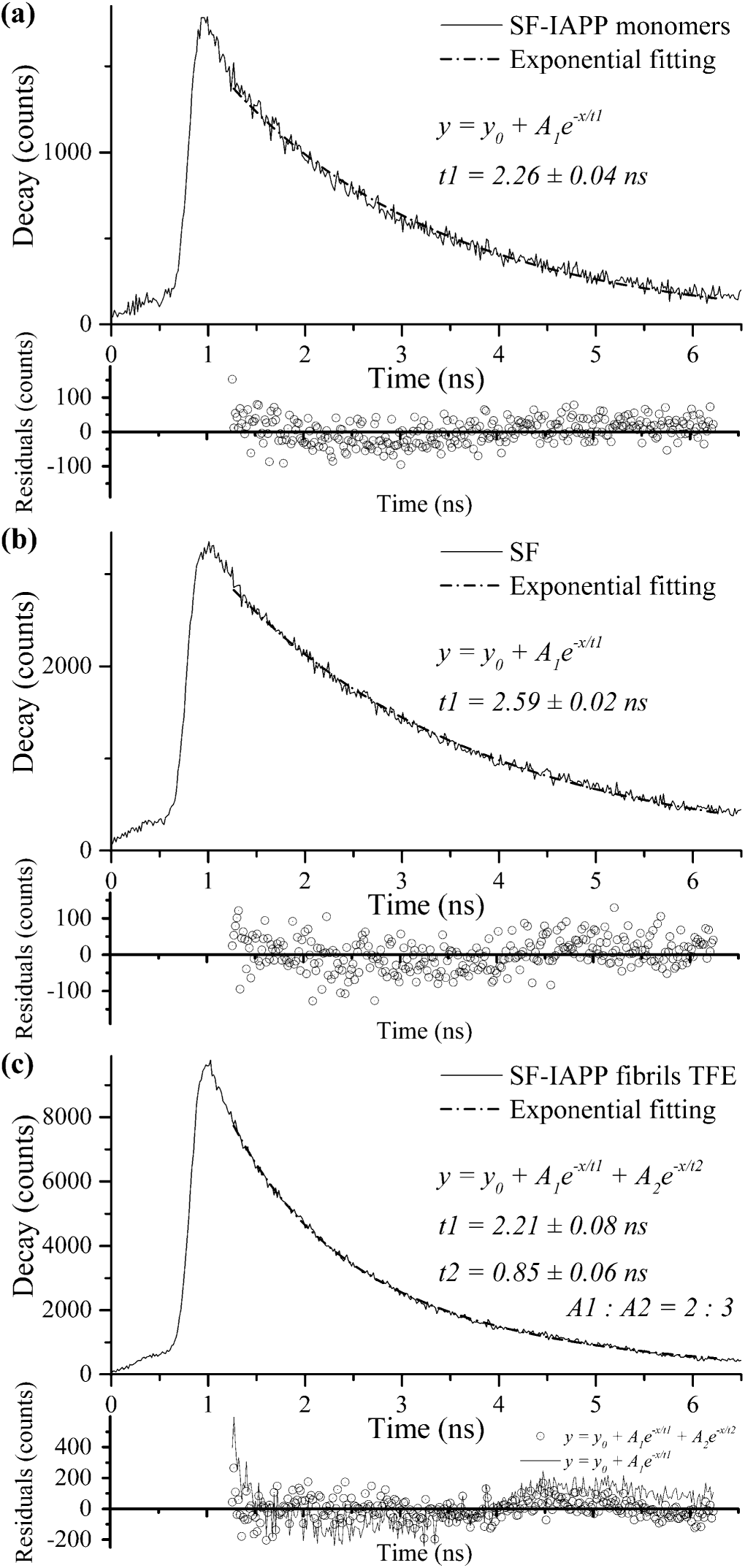
Fluorescence decay and exponential fitting of (a) SF-IAPP monomers, (b) SF monomers, and (с) SF-IAPP fibrils.

Thus, it can be assumed that, during the formation of fibrils, a considerable part of the protein acquires an additional, shorter fluorescence lifetime *t2* (about 0.8 ns) which leads to a total decrease in the average fluorescence lifetime of fibrils relative to monomers in the single-exponential approximation. It should be noted that the time *t1* for monomers and fibrils remains unchanged: approximately 2.2 ns (compare *t1* in Figure 3c and Table 3). Also, the SF lifetime (approximately 2.6 ns) is comparable to one previously reported [12].

The results obtained coincide qualitatively with those described previously for the YFP-alpha-sinuclein fusion protein [6]. This suggests the possibility of using this method for diagnosing the formation of intracellular fibrils *in cellulo* or for the screening of agents that prevent IAPP fibrillogenesis. Thus, on the one hand, the ability of the SF-IAPP fusion protein to form fibrils was demonstrated, yet on the other, the characteristic change in the fluorescence lifetime of the SF protein, as a portion of the fusion protein incorporated into amyloid-like fibrils, was observed. The properties observed likely can be used in methods for detecting the formation of SF-IAPP fibrils *in situ*.

## Conclusion

In its role as part of the SF-IAPP fusion protein, IAPP does not lose the ability to form amyloid-like fibrils. During bacterial SF-IAPP production and subsequent storage, changes to SF-IAPP occur. In particular, single or double amino acid residues undergo deamidation at sites corresponding to the IAPP part. Under the same conditions, SF and SF-IAPP monomers have similar fluorescence time characteristics and the average fluorescence lifetime of SF-IAPP in fibrils decreases.

## Acknowledgments

We thank Mr. Edward Ramsay for help in writing the manuscript.

